# Hearing restoration by a low-weight power-efficient multichannel optogenetic cochlear implant system

**DOI:** 10.1101/2020.05.25.114868

**Authors:** Lukasz Jablonski, Tamas Harczos, Bettina Wolf, Gerhard Hoch, Alexander Dieter, Roland Hessler, Suleman Ayub, Patrick Ruther, Tobias Moser

## Abstract

In case of deafness, electrical cochlear implants (eCIs) bypass dysfunctional or lost hair cells by direct stimulation of the auditory nerve. However, spectral selectivity of eCI sound coding is low as the wide current spread from each electrode activates large sets of neurons along the tonotopic axis. As light can be better confined in space, optical cochlear implants (oCIs) promise to overcome this shortcoming of eCIs. This requires appropriate sound processing and control of multiple microscale emitters. Here, we describe the development, characterisation, and application of a preclinical low-weight and wireless LED-based multichannel oCI system for hearing restoration and its companion to its sister eCI system. The head-worn oCI system enabled deafened rats to perform a locomotion task in response to acoustic stimulation proving the concept of multichannel optogenetic hearing restoration in rodents.

## Introduction

According to the World Health Organization, in 2018 there were 466 million people in the world with hearing loss^1^. It is forecasted that in 2030 this number will reach 630 million and will further grow to 900 million in 2050. If profound, hearing loss makes people potential candidates for hearing rehabilitation by electrical cochlear implants (eCIs). Currently there are approximately 700,000 eCI users worldwide, most of whom achieve good open-set speech understanding in the quiet. eCI systems, associated with low risk for implantation and device failure^2–4^, are composed of an external speech processor and implanted stimulator. They convert sound into electrical current pulses delivered via an intracochlear electrode array to stimulate the spiral ganglion neurons (SGNs), which are ordered according to their characteristic frequency along the tonotopic axis (place-frequency map) that follows the spiral anatomy of the cochlea. Sound processing involves decomposition into frequency bands and extraction of the intensity within each band. These intensities are then used to scale the amplitude of electrical pulses delivered to the electrode at the tonotopic position corresponding to the respective frequency band.

However, due to the wide spread of the electrical current from each of the 12–24 eCI contacts (depending on manufacturer^5^), signals containing information of a given frequency band activate a large fraction of the tonotopically ordered SGNs. This results in limited spectral resolution of sound coding with typically less than ten perceptually independent stimulation channels^6–8^. Poor spectral resolution is commonly considered the bottleneck of the eCI that makes more complex listening tasks, like communication in noisy or reverberant environments, difficult and limits music appreciation^9–11^. Efforts to improve the performance of the eCI include current steering using multipolar stimulation ^12^ as well as intraneural stimulation^13^. Light offers an alternative mode of SGN stimulation with the potential to overcome this bottleneck (reviewed in ref. 14–17). Considering the ability to confine light in space, future optical cochlear implants (oCIs) could activate smaller fractions of SGNs and, hence, enable a higher number of perceptually independent stimulation channels. Two approaches toward optical SGN stimulation have been employed: i) infrared direct neural stimulation (INS)^18^ and ii) optogenetics^19^. While the INS concept has remained controversial for the cochlea^20–23^, optogenetics offers a defined molecular mechanism, restores auditory function in various animal models of deafness and has been successfully implemented in preclinical animal studies by several laboratories (e.g. ref. 24–28). The spectral selectivity of optogenetic SGN stimulation has been shown to be greater than that of electrical stimulation^19,29^ by recordings of midbrain activity and near physiological SGN firing rates can be achieved with fast opsins such as Chronos and f-Chrimson^30,31^.

In parallel, major advances have been achieved towards the technological implementation of the oCI. Since the proof-of-concept study on flexible oCIs based on microscale gallium nitride (GaN) light emitting diodes (μLEDs)^32^ their optimisation (light extraction and focusing) and technical characterisation has been progressed^33,34^. In addition, studies with larger emitters^35,36^ and waveguides^37^ have been undertaken. However, to the best of our knowledge, implementation, characterisation, and application of a sound-driven oCI system has not yet been presented. Such an oCI system should employ real-time sound processing and coding strategies employing more stimulation channels than current eCI. Here, we report the development and functional demonstration of a low-weight, wireless, battery-powered oCI sound processor and driver circuitry to be head-mounted for experiments on freely moving animals. We demonstrate the function of the oCI system and its sister eCI system. In summary, this preclinical oCI system will help to pave the way for developing the future clinical oCI for improved hearing restoration in human deafness.

## Results

### The oCI sound processor and driver circuitry

In order to enable behavioural studies on preclinical optogenetic hearing restoration, we developed, characterised, and applied a miniaturised oCI system consisting of a sound processor, a driver circuitry, and an emitter array. For comparison of eCI and oCI systems, we also implemented a preclinical eCI system. These oCI and eCI systems capture and process sound in real time to drive emitters or electrodes for neural stimulation (Figure 1). Appropriate benchmarking of the oCI system imposes maximum design similarities, i.e. use of the same input stage and digital signal controller (DSC), accommodating differences where appropriate, primarily at the output for driving eCI (charge balanced biphasic current pulses) or oCI (monophasic driving current for light pulses). While the clinical eCI system consists of an external sound processor and an implanted stimulator composed of driver circuitry and electrode array, the miniaturised preclinical oCI and eCI systems combine sound processor and driver circuitry in one battery-powered head-worn assembly (Figure 1 B). The oCI system employed previously described microfabricated LED arrays ^35^ while the eCI system used clinical-style rodent electrode arrays (see Methods). The presented implementation of the oCI driver can employ hundred-twenty-eight stimulation channels of which, due to oCI design, we employed control up to ten LEDs while the eCI driver was designed to operate up to ten stimulation channels of which, due to eCI design, we employed two in the in vivo experiments on rats (see Methods).

**Figure 1.**
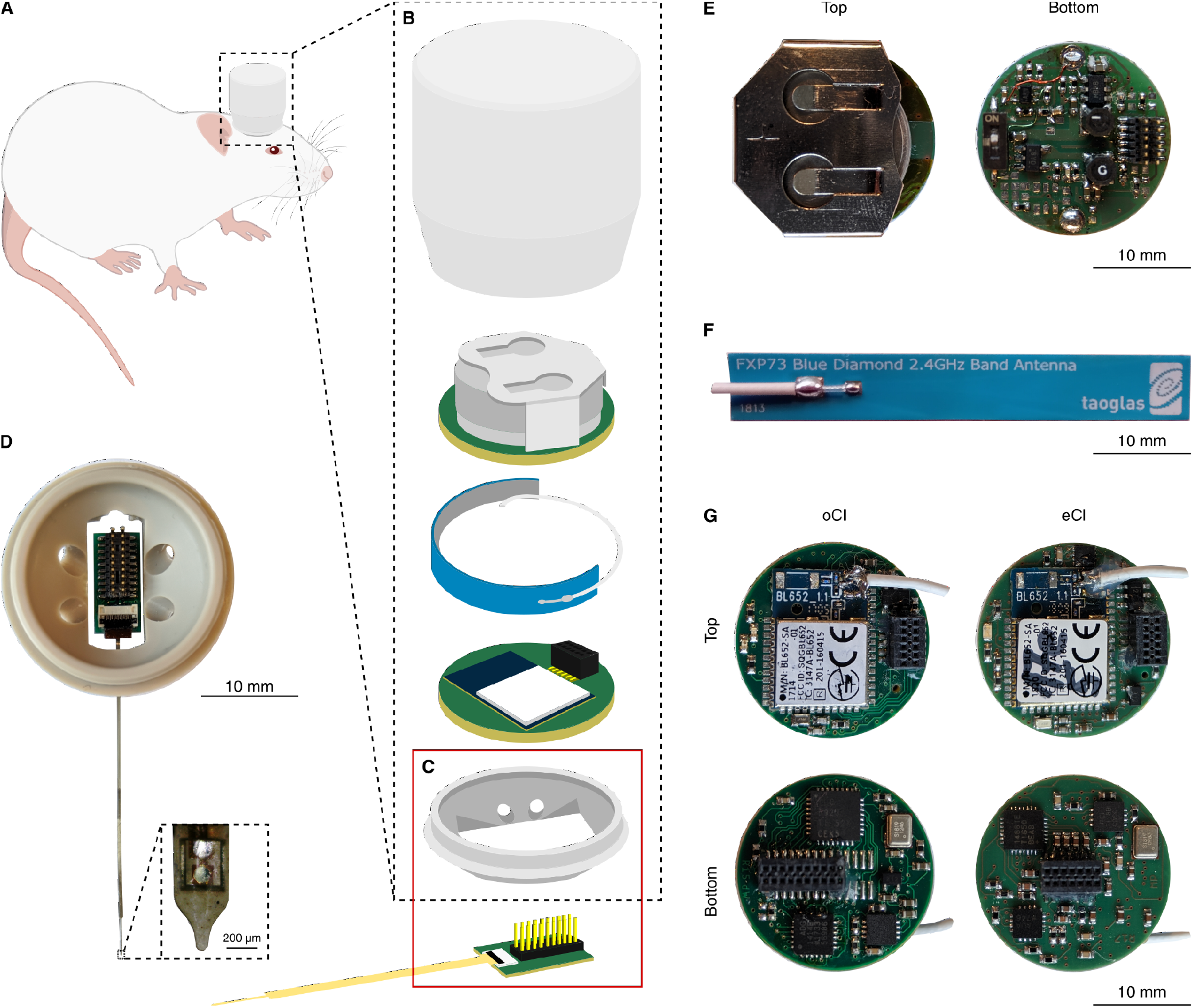
The complete oCI and eCI systems. (A) Illustration of the complete head-mounted oCI and eCI systems. (B) Exploded view illustration of the oCI system consisting of (bottom to top) the oCI with ZIF connector and board interfacing it to driver, head-mounted base of the enclosure, oCI interfacing board, printed circuit board (PCB) containing the sound processor and driver for oCI, wrapped around flexible antenna, powering PCB including rechargeable battery, and screwable enclosure cap (eCI follows the same concept). (C and D) Head-implanted base and connector board for oCI showed in detail in photograph (D, including close-up photograph of the most apical LED of the oCI probe). (E) Photographs of top and bottom view of the powering PCB including rechargeable 3.7 V lithium-ion battery CP1654A3 (VARTA Microbattery GmbH). (F) Photograph of flexible antenna FXP73 (Taoglas). (G) Photographs of top and bottom view of the sound processors and drivers for oCI and eCI.

Size, weight, and portability compatible with head-mounting were among the most important design requirements. The size of the round multilayer rigid printed circuit boards (PCBs) carrying commercial off-the-shelf electronic components measures 20 mm in diameter (Figure 1 E and G). The stack of two PCBs integrating all components including batteries measures 20 mm in height. The whole assembly is housed in a light and robust enclosure consisting of a base (mounted to the animal’s skull) and a screwable cap for convenient battery replacement. The weight of the complete sound processor including enclos-ure and batteries is below 15 g, which is around 33 % of the animal’s head weight (40–45 g). Minimizing the energy consumption of the sound processor was another design criterion with particular relevance for the choice of a digital signal controller (DSC) and the implementation of wireless communication. For system details and choice of DSC see Supplementary Information.

The chosen DSC (nRF52832, Nordic Semiconductor) connects to a computer (PC) wirelessly over its 2.4 GHz radio transceiver using a protocol based on Enhanced ShockBurst (ESB), or over a 2-wire universal asynchronous receiver-transmitter (UART) port (Figure 2). Further inputs to the DSC are the PDM microphone, an operational amplifier for electrophysiological recordings via its analogue-to-digital converter (ADC), and a trigger input via a general-purpose input/output pin using tasks and events (GPIOTE). The battery board (not illustrated in Figure 2) is also connected internally to the ADC to be able to monitor battery voltage. For output, the driver circuitry of either oCI or eCI is connected via two SPI instances.

**Figure 2.**
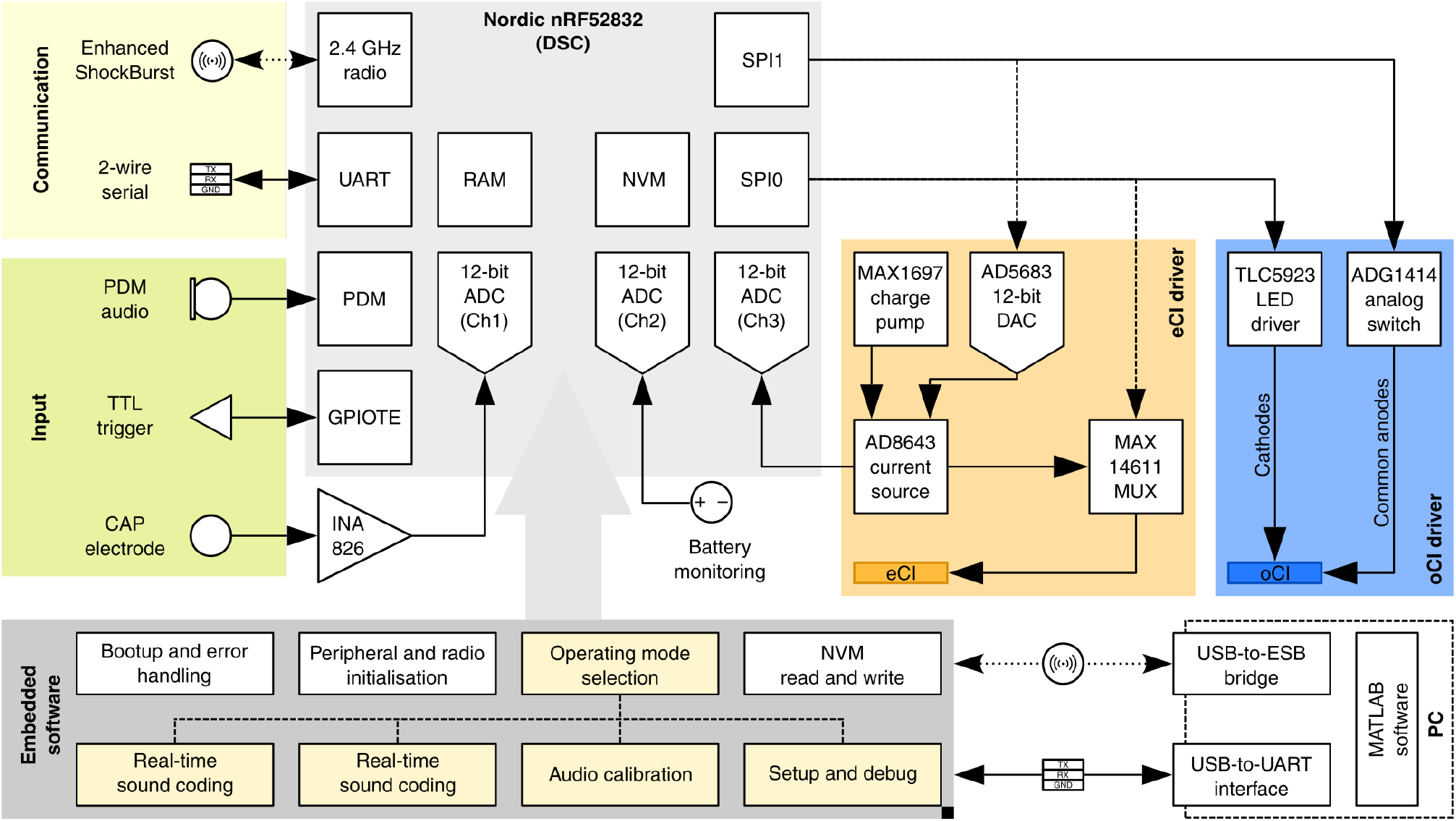
Overview of the oCI and eCI systems. Signal pathways are schematically represented by arrows, the orientation of the arrowheads indicate the signal direction. Except for the 2.4 GHz radio transceiver, every arrow denotes wired connection. The communication block (yellow) indicates wired (UART, universal asynchronous receiver-transmitter) and wireless (ESB, Enhanced ShockBurst) communication interfaces. The input block (green) uses the Nordic nRF52832 digital signal controller’s (DSC’s, light grey) pulse density modulation (PDM) interface for audio input, one analogue-to-digital converter (ADC) channel for electrophysiological recording, one ADC channel for monitoring the battery, and a general-purpose input/ output pin using tasks and events (GPIOTE) for high-precision triggering. The serial peripheral interface (SPI) ports of the DSC are connected either to the oCI driver circuitry (blue: LED driver, analogue switch, and oCI interface) or to the eCI driver circuitry (orange: current source, multiplexer (MUX), a digital-to-analogue converter (DAC), and eCI interface). In the eCI case the third ADC channel is used for the voltage feedback control loop of the output of the current source enabling electrode impedance measurement. The oCI and eCI systems are powered by a rechargeable 3.7 V battery. Embedded software (dark grey) handling different operation modes runs on the digital signal controller. The software framework boots up the device, initialises peripherals connecting external components, and saves settings and logs to the non-volatile memory (NVM). Limited random-access memory (RAM) imposes the software framework switching dynamically between modes of operation in which related memory allocations and deallocations as well as starting and stopping of peripherals are done. Setting up and debugging the device is possible over wireless (2.4 GHz) or wired communication interfaces.

In the oCI system, the DSC controls a 16-channel digitally-adjustable current sink (TLC5923, Texas Instruments Inc.) and an 8-channel analogue switch (ADG1414, Analog Devices) via its SPI0 and SPI1 interfaces, respectively. Such configuration enables up to 128 channels in a matrix addressing concept^32^. While the analogue switch can provide voltage eslectively to common anodes of LED blocks, the current sink can adjust the exact amount of current (hence light emission) of each LED within a block. It takes around 15 μs for a DSC-internal command to set up the complete LED array (independent of the topology of the LED array) with the requested currents. In the eCI system, the DSC’s SPI ports control a 16-channel multiplexer (MUX; MAX14661, Maxim Integrated) and a 12-bit digital-to-analogue converter (DAC;AD5683, Analog Devices) driving a current source (AD8643, Analog Devices) to select the electrode contacts to be actuated and to supply the charge balanced biphasic stimuli, respectively. The current source voltage can be monitored by the DSC via an ADC channel. The oCI and eCI driver provides the output via pin headers interfaced with the stimulation probes (for details see Methods and Supplementary Information).

For the audio input to the oCI/eCI system, we used a microelectromechanical system (MEMS) microphone (SPH0641LM4H-1, Knowles) interfaced to the DSC via pulse density modulation (PDM) interface providing sufficient sensitivity to sound frequencies at least up to 25 kHz (Supplementary Figure 3) in order to cover a substantial part of the audible spectrum of rats (1–70 kHz)^38,39^ and marmoset monkeys (0.1–35 kHz)^40,41^. Also the selected microphone is energy efficient (~0.8 mA current consumption during use at 3.3 V) and has both a wide dynamic range as well as a relatively flat transfer characteristic (Supplementary Figure 3). For both eCI and oCI systems the battery enabled 7–8 hours of operation when driven with identical conditions as used in behavioural experiments (typical session duration is less than an hour; for details see Methods and Supplementary Information).

### Software framework

For evaluating the preclinical oCI system in comparison to the eCI, the software framework considers several functionalities that encompass a subset of features of clinical CIs (Figure 2). Real-time sound processing (used for behavioural assessment) and triggered stimulation protocols (for physiological and behavioural assessment) comprise the two main modes of operation. The software framework switches dynamically between operating modes and manages related memory allocations and deallocations as well as starting and stopping of peripherals. This concept was introduced for three reasons: (1) the limited random-access memory (RAM) does not allow the DSC to cater to all functionalities at the same time, (2) turning off peripherals not relevant to a given task saves energy hence extending battery life, (3) task-specific grouping of functions contributes to safe operation and robustness against unintended use.

In addition, the software framework encompasses standard functions such as booting the device, initialising peripherals, and saving settings and logs to the non-volatile memory (NVM). It also provides managed access to 4 kB of the NVM, where initial parameters (among which is a unique identifier) are burned-in before first operation of the DSC. These parameters can be changed and overwritten later during operation in “Setup and debug” mode (Figure 2) via specific commands over wired or wireless communication channels. The NVM is also shared with a simple embedded logger that can store critical logs for later read-out.

The software operates on an ARM Cortex-M4F central processing unit (CPU)—a reduced instruction set computer (RISC) at the heart of the nRF52832 DSC—featuring three-stage instruction pipeline, Thumb-2, digital signal processing (DSP), and single-precision floating point instructions at excellent energy efficiency^42^. Thanks to the DSP library of the Cortex Microcontroller Software Interface Standard, a suite of common signal processing functions is available as highly optimised code for the platform^43^.

### Sound processing and spectral decomposition

Whenever the signal processor is in real-time sound encoding mode, it transfers audio samples from the PDM microphone to the RAM, spectrally decomposes the signal, and transforms it into patterns of activation of the oCI emitters or the eCI electrodes aiming to stimulate SGNs at the tonotopic positions that correspond to the frequencies contained in the sound signal. While sound capturing—once appropriately configured and started—operates by direct memory access and double buffering without substantial involvement of the CPU, the further processing of the audio samples places major demands on the computational power. For this reason, we carefully evaluated spectral decomposition via FFT -vs. IIR-based filter bank operations (Supplementary Figure 1).

Because the optical or electrical stimulation channels are placed equidistantly on the cochlear implant, the employed filter bank should ideally have a quasi-logarithmic frequency resolution for the stimulation to match the tonotopic organisation of the cochlea. This is easily set up when using an IIR-based filter bank: with IIR filters the required computational power scales almost linearly with the number of required frequency channels. When using FFT, the increase of computational power as a function of frequency channels is less obvious: the real FFT implementation of the DSP library is very efficient but it implies various overheads in our case. First of all, every block of input data needs to be multiplied by the values of a window function. Then, real-valued magnitudes need to be calculated from the complex spectral values, requiring not only multiplications and additions, but also the computationally more expensive square root calculation. Next, the resulting magnitudes, corresponding to the linearly spaced real FFT frequency bins, need to be combined to form quasi log-spaced channel frequencies, which requires further multiplications and additions. Finally, processing encompasses 50 % overlapping of the temporal analysis windows to not miss transients, which, as a consequence, doubles the required calculations. For these reasons, we chose an IIR-based filter bank for spectral decomposition. More specifically, we used the single precision floating point implementation of a biquadratic second order IIR (biquad) band-pass filter as the basic building block of our filter bank with quality factor of 12. We also evaluated the fixed-point implementation but found only a trend toward better performance at the same current consumption, while trading in a reduced dynamic range and noise artefacts due to rounding errors.

### Sound coding strategy

The efforts of optogenetic hearing restoration build on the hypothesis that optical sound encoding has near-physiological spectral selectivity^29^. Following the Nyquist sampling theorem, the number of independent optical stimulation channels should be at least twice the number of the critical bands the cochlea holds. This would equate to at least 48 channels for the human cochlea^44^ and ~8–12 for the rat^45^. At the same time, the most efficient pulse duration for optogenetic stimulation which depends on the kinetics of the opsin of choice (0.5–2 ms^24,30,31^) exceeds that of electrical stimulation (typically below 100 μs^46^). Moreover, the synchrony of optogenetically driven SGN firing^24,30,31^ is lower than that of electrical stimulation^47–49^ and closer to physiological sound stimulation^24,30,31^. This indicates a choice of ~1 ms optical stimulation and limits the maximal stimulation rate to SGN-typical steady state firing rates of 300 Hz^50–54^.

Here, we implemented the continuous-interleaved sampling (CIS) strategy ^55^ and the *n*-of-*m* strategy^56^, two state-of-the-art clinically used coding strategies for the sake of a first preclinical optical coding strategy as well as for operating the eCI. In the case of the *n*-of-*m* strategy, *m* denotes the total number of addressable stimulation sites of the implant, whereas *n* is the maximum number of stimulation sites to be used in a stimulation cycle. Typically, one stimulation cycle is based on the spectral analysis of one buffer of input audio samples and runs sequentially through the stimulation sites from the base to apex once while bypassing those considered least influential with usually *n* ≤ *m*/2 (ref. 56).

In our implementation, real-time sound processing encompasses three concurrent software processes. The first is maintaining sample transfer from the PDM microphone to a set of buffers, the size of which depends on the required spectral update rate. Whenever a buffer got filled up with new audio samples, the second process performs the core calculations of the coding strategy and stores the order and corresponding magnitude of stimulations for a complete stimulation cycle into the stimulation buffer. The third process is launched periodically at the rate of stimulation of the oCI/eCI, fetches stimulation parameters from the buffer, and orchestrates hardware peripherals to output the upcoming stimulus with high temporal precision.

The core calculations of the sound coding strategy process start by spectrally decomposing the signal into *m* bands using a bank of second order IIR filters (as detailed in Sound processing and spectral decomposition) and extracting the envelopes of the band-filtered signals. Next, either all spectral values (CIS, where *n* = *m*) or just the *n* main spectral values (*n*-of-*m*) are taken and converted into dB relative to full scale (dB FS) values. The latter is denoted *x*_*i*_ for channel *i*. Next, *x*_*i*_ is mapped to the adequate physical stimulation range yielding stimulation amplitude *y*_*i*_ for each channel *i* according to:

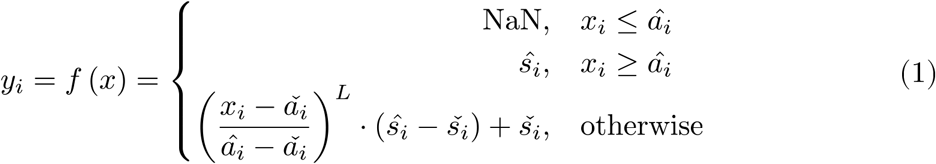

where 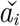 and 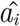 denote acoustical threshold and saturation levels (in dB FS), respectively, 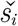 and 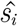 denote threshold and maximum comfortable levels in terms of stimulation intensity, respectively, and *L* denotes loudness growth coefficient, a parameter used to change the non-linearity of the mapping between soft and loud signal parts and is used in human CI systems to tune the perceived loudness growth. While optimal 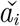 and 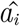 values mostly de-pend on the audio recording hardware (see Supplementary Information), all above parameters are influenced by the acoustic environment the sound processor is intended to be used in. Calculated *y*_*i*_ values can be directly used to set the optical or electrical stimulation circuitry’s magnitude for each stimulation pulse. In case of *y*_*i*_ = NaN, no stimulation will take place on the corresponding channel i in the actual stimulation cycle.

### Wireless control and communication protocol

While the standardisation might be considered as an advantage of Bluetooth Low Energy (BLE)—e.g. offering remote controlled by any BLE-capable device—we anticipated substantial drawbacks. BLE requires larger overhead (in terms of code space, working memory that must be dedicated to its functionality, and other resources e.g. timers) than a proprietary radio protocol. Furthermore, the additional protocol stack layers of BLE cause a latency in transmitting a single short message in the range of 10 ms (measured from enqueueing the message on the sender side till receiving the message contents on the receiver side, see also ref. 57). Since short trigger latency was one of the key design requirements, we choose to build upon Nordic Semiconductor’s proprietary ESB protocol. ESB avoids most of the BLE-typical overhead thereby freeing up peripherals, code space, and working memory of the DSC, also enhancing wireless responsiveness. The use of the ESB allowed to achieve the time between sending a wireless trigger command and a stimulation start in the range of approximately 350 μs. Disadvantages of ESB include a shortage of sophisticated 2.4 GHz wireless technologies (manual frequency channel selection required) and the need for ESB-capable communication interface on the PC side. For this reason, we implemented a custom-coded firmware running on a single-board development kit for 2.4 GHz proprietary applications (nRF52 DK, Nordic Semiconductor) connected to the PC via a USB (USB-to-ESB bridge). To increase speed of data transmission, commands sent from a PC via USB interface and arriving at the bridge are not interpreted by its firmware but forwarded directly to the oCI/eCI system. A TTL trigger input can be provided to the GPIOTE of the USB-to-ESB bridge and sent as a triggering command wirelessly to oCI/ eCI system or, in a wired configuration, directly to the GPIOTE of the oCI/eCI system. Both wireless and wired communication protocols also allow triggering via a command sent from the PC. For the radio operation of the USB-to-ESB bridge and oCI/eCI system, different modes of radio operation were implemented in each firmware. Permanent receiving or transmitting of radio is always turned on, for applications where only minimal latencies are acceptable. Periodic radio (transmission turned on with a given frequency) is used for applications with lower requirement for timing but emphasis on low power consumption). Preselection of radio modes for the via USB-to-ESB bridge and oCI/eCI allows for reliable communication and triggering of a predefined stimulation pattern.

The communication protocol (also over wired UART interface) uses fixed-length commands in a form of a human-readable 15-character-long ASCII messages. Due to the lower overhead, interpretation of such messages in real-time and debugging are straight forward. All commands sent to the DSC of the oCI/eCI system (e.g. application of new setting) can be acknowledged to ensure success of communication or retrieve requested information (e.g. status of the DSC or its settings). The configuration is stored in the NVM and loaded at boot. The performance of the wireless communication of a PC with a processor via USB-to-ESB bridge was assessed in the animal experiments (see In vivo experiments).

### In vivo experiments

In order to test whether optical/electrical stimulation with our oCI/eCI sound processor triggers behavioural responses, we turned to instrumental conditioning by negative reinforcement (avoidance behaviour) paradigm in a custom-built ShuttleBox setup (Figure 3 B, for details see Methods). oCI systems were tested on kanamycin-deafened rats, which had received postnatal injection of AAV-PHP.B^58^ carrying the ChR2 mutant CatCh^59^ under the synapsin promoter into the left cochlea. Four adult rats were implanted with unilateral (left side) chronic oCI probes. For comparison, additional kanamycin-deafened rats were implanted with eCI probes (4–8 month of age, N = 6 females; for details on surgical procedures see Methods).

**Figure 3.**
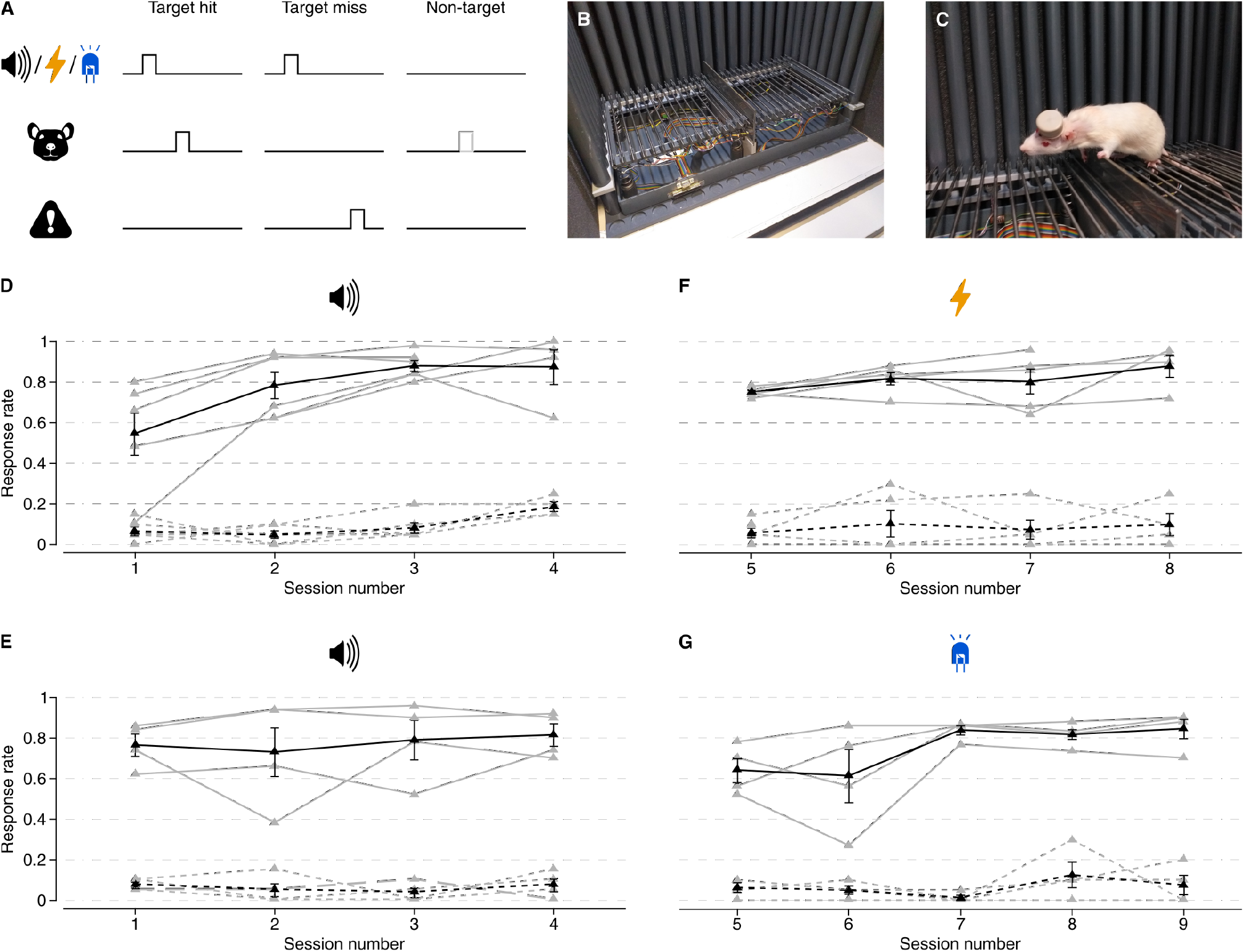
Behavioural responses to electrical (eCI) and optical (oCI) stimulation of the cochlea. (A) Outline of the ShuttleBox paradigm. Upon a sensory cue (acoustic click, electrical or optical) in the target trials, the animal needs to cross the hurdle, otherwise an electrodermal stimulation is delivered via the grid floor of the setup. If the animal crosses the hurdle in time, the trial is considered successful (left), otherwise the trial is considered a fail (centre). Non-target trial without cue and behavioural consequence are intermingled to assess the baseline activity of the animal (right). (B) Image of the ShuttleBox setup, placed in a sound attenuating chamber. The doors of the sound attenuating chamber are open and the front wall of the ShuttleBox is removed for illustration purposes. (C) Image of a trained rat, carrying head-mounted oCI system, performing in the ShuttleBox. (D and E) Response rates to a target (solid lines) and non-target (dashed lines): acoustic stimulation using 70 dB clicks prior eCI (D, N = 6) and oCI (E, N = 4) implantation; (F and G) predefined stimulation delivered by the sound processor via both eCI electrodes at intensities between 100– 200 μA (F, N = 6) and all oCI LEDs at intensities between 10–100 % (G, = 4). Black symbols and lines indicate mean of data ± SEM.

Before implantation animals (both eCI and oCI groups) were trained with acoustic click stimulation in the ShuttleBox paradigm (for details see Methods) to achieve response rate of at least 0.8 for a target while not exceeding 0.2 for non-target control trial (Figure 3 D and E). After deafening, implantation, and adaptation of the animals to the new stimuli, animals performed the same task as previously, reaching similar rates for predefined stimulation with both eCI (Figure 3 F) and oCI (Figure 3 G). In both cases all available stimulation contacts were used, i.e. both eCI electrodes at intensities between 100–200 μA (mean threshold was ~70 μA) for eCIs and all oCI LEDs at intensities between 10–100 % (~0.31–3.1 mA LED current per channel).

After we had shown that the predefined stimulus from the eCI and oCI systems can elicit behavioural response (Figure 3), we tested if real-time sound coding by oCI system can restore hearing in deaf (Figure 4) and partially deafened animals (Supplementary Figure 4). Hearing was evaluated before and after deafening by recording acoustic auditory brain stem responses (aABRs) as well as by the ShuttleBox paradigm (Figure 4, Supplementary Figure 5 A–C). Using all LEDs inside the cochlea we aimed to restore optically sound coding using both, *n*-of-*m* and CIS coding strategies, implemented into the firmware for *m* = 8 (Figure 4); *m* = 6 (Supplementary Figure 5) and *n* = 3 (in case of *n*-of-*m*; for details see Sound coding strategy). We roughly estimated threshold level (TL), the most comfortable level (MCL), and a microphone gain as we had observed that operating all LEDs at intensity levels between 30–100 % triggered behavioural responses but no aversive behaviour like freezing or intensive scratching in the animals (see also Methods). We evaluated the avoidance behaviour in the ShuttleBox in response to acoustic stimulation in a deaf rat with a fitted oCI system (Figure 4 C and D). Initially, we employed wideband noise stimuli (frequency ranges: 2–10.5 kHz and 10.5–20 kHz) activating up to 3 LEDs and triggering the avoidance behaviour. We then moved on to narrowband noise bands recruiting fewer LEDs and, again, could demonstrate an auditory percept using oCI sound coding. In another animal which was only partially deafened (high frequency hearing loss), the sound-driven oCI-system elicited comparable response rates for impaired and non-impaired frequency ranges (Supplementary Figure 5 D and E). The operation of the oCI system was strictly required for the behavioural response: sound did not elicit the behaviour when the oCI was switched of (Figure 4 C and D, black line; Supplementary Figure 5 C).

**Figure 4.**
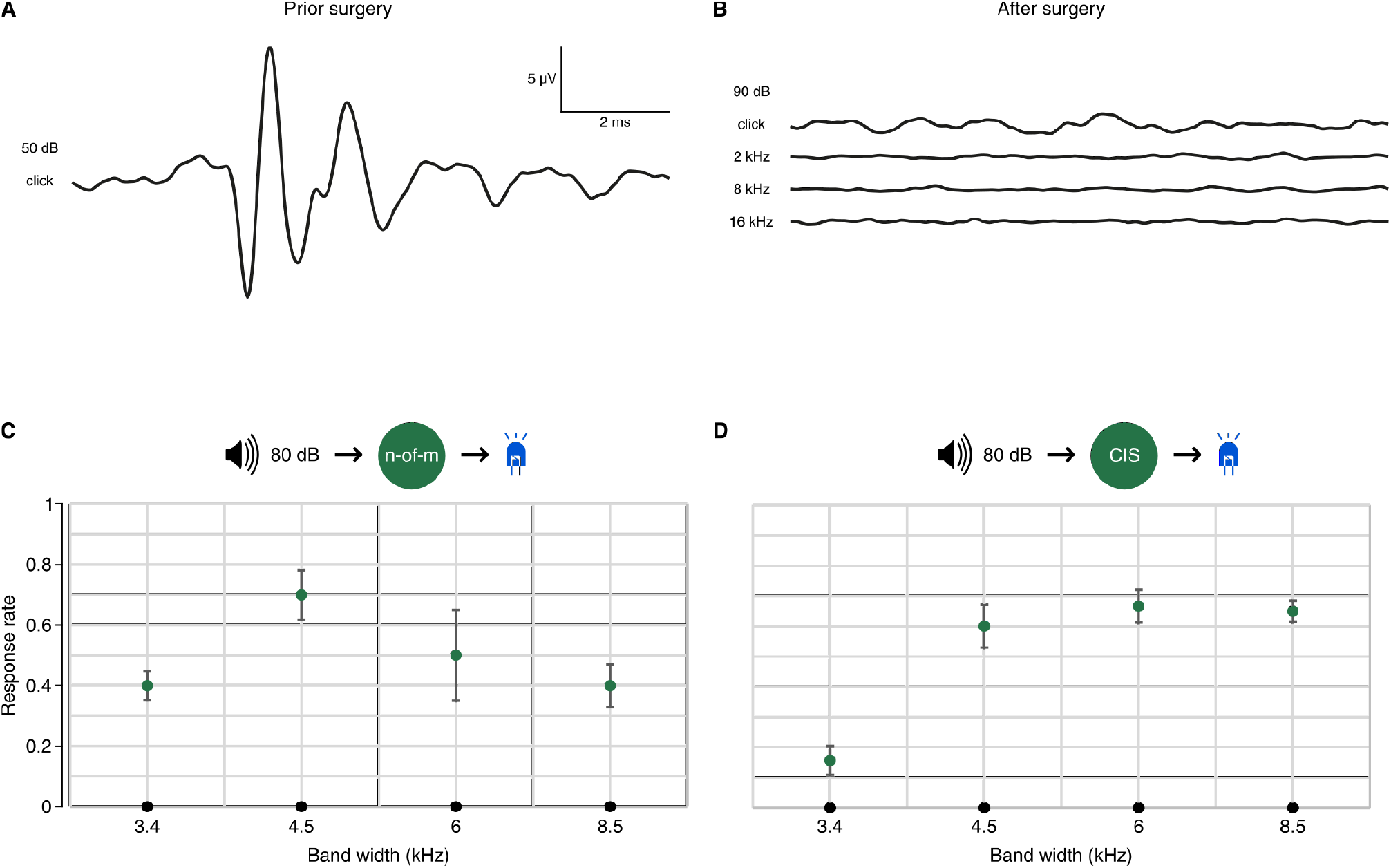
Restoring hearing with the oCI system using different coding strategies of the sound processor. (A) Representative aABR, in response to clicks at 50 dB prior implantation and deafening. (B) aABRs of the same animal after oCI implantation and deafening in response to 2, 8, and 16 kHz clicks at 90 dB. (C and D) Results of the ShuttleBox sessions with activated oCI system encoding noise bursts of different band widths spanning the range of 2–20 kHz into a patterned optical stimulation via 6 LEDs inside the cochlea of the AAV-transduced rat using either *n*-of-*m* (C) or CIS (D) coding strategy in a deafened rat. Within single sessions different widths of frequency bands covering the 2–20 kHz range were presented to the animal as target trials (3.4 kHz: 5 bands, 4.5 kHz: 4 bands; 6 kHz: 3 bands; 8.5 kHz: 2 bands). In each session, additional non-target trials were presented. Response rate is a response to the target trials subtracted by response to non-target trials. Green dots represent mean response rate (± SEM) to all ranges within each session. Black dots represent control session in which the oCI system was not activated: no response was observed. All acoustic stimuli were presented at 80 dB.

## Discussion

Here we describe, characterise, and apply a miniaturised multichannel oCI system for animal studies and compare it to its sister eCI system. Key features include low weight, realtime sound processing for multichannel stimulation (oCI: up to 128 channels in the matrix configuration, eCI: 10 channels), wired or wireless control, and operation over several hours in behavioural experiments. In order to enable rigorous benchmarking of the oCI, the design of hard- and software for oCI and eCI was kept as similar as possible, such that the two systems primarily differ in the number of stimulation channels and precise driver hardware. Next, to technical characterisation and benchmarking of components, we provide a proof-of-concept of the oCI-system using optogenetically modified rats. This study proves the feasibility of using the miniaturised oCI system to restore auditory-driven behaviour in deaf, freely moving rats.

### Utility of the developed oCI system

The oCI sound processor and driver circuitry operated “active” oCIs^60^ based on microfabricated 1D-arrays of ten blue GaN LEDs for multichannel optogenetic stimulation of the auditory pathway. However, the probes used here did not fully capitalise on the potential of the oCI sound processor and driver circuitry (e.g. real-time 32 channel sound processing) given the small size of the rat cochlea. The developed oCI systems pave the way for further preclinical development of multichannel optogenetic stimulation of the auditory pathway in animal models with larger cochlea such as gerbils, cats or marmoset monkeys^61^. Moreover, the hard- and software framework established here can as well drive more sophisticated arrays of emitters also for other optogenetic applications in basic and preclinical research. For instance, using the implemented matrix addressing^32^ it can operate 2D arrays or more complex 1D arrays, in which emitters of one block share a common cathode. This is highly relevant for when scaling-up the number of channels in “active” oCIs^60^ e.g. based on μLED arrays^32,33^ where the number of lines is limited by space constraints of the scala tympani. The same sound processor and a similar driver circuitry can also serve for the control of laser diodes in multichannel “passive” oCIs^60^ where the light is fed into the cochlea via polymer-based waveguides arrays^62^. Future modifications of the described hard- and software framework will also accommodate the combination of electrical and optical stimulation (combined oCI and eCI) for hybrid electro-optical stimulation^26^ or optofluidics circuitry for application of photopharmacology into the cochlea and later optical stimulation^63–65^. Most studies presenting optogenetic stimulators address brain stimulation^66–69^, e.g. cortical control of movement^70^, and also peripheral nervous system^71^. Some systems require external devices to deliver stimuli^72^: via wireless transfer of power (inductive coupling) from an external transmitting coil to a device coil or radiofrequency power harvesting from an external transmitter antenna to a receiver antenna located on the device. Some of these devices, similarly to ours, enable wireless communication and battery powering, being even lighter. However, our system not only can stimulate with predefined patterns set via a wireless communication protocol but can process captured sound in a real-time and convert it into optical stimulation pulses, thereby operating as a fully stand-alone system similar to clinical stat-of-the-art eCI systems.

### Towards clinical translation of the oCI system

In order to implement a basic, head-wearable multichannel oCI system for animal experiments we emphasised low-weight and small size. The weight of the whole system corresponded to roughly one-third of the animals head weight, which is well below the provisions of the U.S. Army Aeromedical Laboratory of 50 % (see also ref. 73). Moreover, a study in rats reported no adverse effects in animals carrying nearly twice as much weight^74^ as we used here. Different from clinical eCI systems consisting of an external sound processor and the actual implant with driver and electrode array, we implemented a single, external head-mounted device to which the oCI or eCI probe were interfaced at the vertex of the animal’s skull. However, our approach shows feasibility of driving oCI with a miniaturised system built with commercial off-the-shelf components and as a proof of concept, paves the way for the future development of human prototypes based on dedicated components.

This will closely follow the design of current eCI systems but will accommodate the specific features of the oCI such as matrix addressing of oCI such as matrix addressing^32^ of a greater number of stimulation channels. Challenges en route to clinical translation of the oCI include the co-development and application of cochlear gene therapy as well as optical stimulation introducing new technology into the CI and potentially having a greater power budget and being phototoxic. Given the higher per pulse energy requirement of oCIs with currently available channelrhodopsins^17^ and the larger number of stimulation channels, saving energy on other ends will be critical. For instance, the stimulation rate can likely be reduced from (~1 kHz) to 300 Hz close to maximal physiological steady state firing rates of SGNs or even adapted to stimulus intensity to mimic physiological extend the range of rate coding. A lower rate of optical stimulation seems justified by speech recognition with eCI sound coding not improving much beyond stimulation rates of 500 Hz^75^. Moreover, the greater stochasticity of optogenetically induced firing suggests that high stimulation rates (~1 kHz) as employed in eCIs to invoke SGN refractoriness for avoiding non-natural SGN synchronicity^46^ will not be required in oCIs. Dedicated optical coding strategy will also need to consider the design of the oCI. For “active” oCIs, i.e. active optoelectronic light emitters implanted into the cochlea^60,76,77^, two concepts have been put forward in order to cope with addressing dozens of emitters while not scaling up the number of feeding lines given the rigid space constraints of the cochlea. The matrix addressing concept^32^ relies on ultrafast activation of the channelrhodopsins (μs range) and integration of the resulting depolarizing current by SGNs for spike generation^78^. A second concept foresees an array of emitters on CMOS that connect by a limited number of lines for operation^79^. In conclusion, to the best of our knowledge, this is the first multichannel oCI system allowing experiments in freely moving animals, paving the way for developing future clinical oCI systems for improved hearing restoration.

## Methods

### Surgical procedures

CI surgeries were performed on animals anaesthetised with isoflurane (3.5–4 % at 1 l/min for induction, 0.5–2 % at 0.4 l/min for maintenance). Appropriate analgesia was achieved by subcutaneous injection of buprenorphine (0.1 mg/kg bodyweight) and carprofen (5 mg/kg bodyweight) 30 minutes prior to surgery. Depth of anaesthesia was monitored regularly by the absence of reflexes (hind limb withdrawal) and breathing rate and adjusted accordingly. Animals were kept on a heating pad to maintain body temperature at ~37°C. All experimental procedures are in compliance with national animal care guidelines and were approved by the local animal welfare committee of the University Medical Center Göttingen as well as the animal welfare office of the state of Lower Saxony, Germany.

Animals were chronically implanted unilaterally (left side) with an eCI or with an oCI into the scala tympani via a cochleostomy of the basal turn. oCI surgery was preceded by an early postnatal injection of AAV-PHP.B carrying the ChR2 mutant CatCh under the synapsin promoter into the left cochlea (at postnatal day 6, 3 –6 months before the implantation). Animals were deafened prior to oCI/eCI implantation: 2–3 μl of kanamycin solution (100 mg/ml; Kanamysel, Selectavet) were injected in both ears. In the left cochlea kanamycin was injected via the cochleostomy which was also used for the implant later, while on the right ear kanamycin was injected via the cochleostomy which was induced via a transtympanic approach.

### SGN stimulation: probes and protocols

Multichannel oCI and eCI probes were designed to fit the dimensions of the scala tympani of the rat cochlea (length of ~7 mm, ref. 80) and have been described in previous studies: eCI^81^ and oCI^82^. They will be briefly introduced.

#### eCI probes

The most apical electrode was positioned ~3 mm-deep inside the cochlea and the return electrode was inserted between connective tissue and muscles in the neck of the animal. The eCI probes consisted of 2 linearly-arranged intracochlear platinum sheet contacts and an extracochlear platinum-iridium reference ball electrode (Supplementary Figure 3 A and B). All contacts were connected via lead wires to a 3-pin male connector (2 mm pitch) and are embedded into silicon. The intracochlear contacts were 0.3 mm in diameter each with a centre-to-centre pitch of 1 mm. To assure reproducibility of implantations across animals an array was marked with a black dot at a distance of 3 mm measured from the apical tip (Supplementary Figure 3 B). The diameter of the silicone-encapsulated intracochlear and intrabullar part was 0.3 mm increasing to 0.9 mm at the extrabullar part to provide a stable submuscular connection. The silicone-encapsulated return electrode was 0.3 mm in diameter. These animal eCIs, except the size and the number of stimulation sites, are virtually identical to implants used in human patients. Monopolar electrical stimuli (charge balanced biphasic cathodic-first pulses) were delivered.

#### oCI probes

oCI contained 10 LEDs (C460TR2227-S2100, Cree), each 220 μm by 270 μm by 50 μm, spaced at a pitch of 350 or 450 μm along a polyimide carrier comprising wiring and LED contact pads encapsulated into silicone (Supplementary Figure 3 C). Its ZIF connector (5020781362, Molex) was interfaced with the head-mounted adaptor board to male pin header (FTE-110-03-G-DV, Samtec) that in turn was connected to the female pin header (CLE-110-01-G-DV, Samtec) of the driver circuitry. Light pulses were delivered via custom-made oCIs driven by monophasic current. oCI fabrication processes have been adapted to the chosen device layout following ref. 35 and ref. 82.

#### ABR recordings

ABRs were recorded using needle electrodes on the vertex and mastoid, while an active shielding electrode on the neck was used to reduce the noise level. The differential potential between vertex and mastoid subdermal needles was amplified using a custom-designed amplifier (gain 10,000), sampled at a rate of 50 kHz (NI PCI-6229, National Instruments), and filtered off-line (0.3 kHz to 3 kHz Butterworth filter) for acoustically evoked ABRs.

Stimulus generation and presentation, and data acquisition were realised using custom-coded MATLAB (The MathWorks, Inc.) scripts operating digital-to-analogue converters (National Instruments) in combination with custom-built hardware to amplify and attenuate audio signals. For acoustically evoked ABRs, sounds were presented in an open near field via a single loudspeaker (Vifa, Avisoft Bioacoustics) placed on average 30 cm in front of the animal at the level of the animal’s head. Sound pressure levels were calibrated with a 0.25-inch microphone (D 4039, Brüel & Kjaer GmbH) and a measurement amplifier (2610, Brüel & Kjaer GmbH).

### Behavioural experiment setup

The setup was identical as previously used in studies of optogenetic stimulation via optical fibre involving Mongolian gerbils^24^. Briefly, the ShuttleBox is composed of two platforms and a hurdle between them. Platforms, consisting of equally spaced rows of round metal bars, are mounted on springs and accelerometers to determine the position of the animal. Each acoustic stimulus is presented through a ceiling-mounted loudspeaker (Vifa, Avisoft Bioacoustics). To generate band noise, white noise was bandpass filtered for specific frequencies and noise bands using built-in functions of MATLAB (The MathWorks, Inc) and verified by spectrogram analysis performed with oscilloscope (DSO-1102D, Voltcraft) running FFT algorithm in combination with a 0.25-inch microphone (D 4039, Brüel & Kjaer GmbH) and a measurement amplifier (2610, Brüel & Kjaer GmbH). Sound pressure levels for each stimulus were calibrated using the same system. Stimulus generation as well as data recording is performed with custom-coded MATLAB (The MathWorks, Inc) scripts via digital-to-analogue converters (National Instruments) and custom-built hardware (for details see ref. 24). The existing custom-coded MATLAB (The MathWorks, Inc) scripts were extended to interface with the oCI/eCI processor for adjusting settings and for linking the oCI/eCI processor with a PC to trigger predefined stimuli (Figure 3).

### Behavioural experiments paradigm

The behavioural training procedure was similar to previously published^24^. First, rats were behaviourally trained in acoustic click sessions, later followed by trainings with oCI/eCI stimulation (with triggered predefined stimuli and sound coding-based acoustic stimulation). For each training session, the animal was positioned on one of the platforms inside the ShuttleBox and adapted for 5 minutes before any stimulation. Per session, target trials were presented with an inter-trial-interval randomised between 12 and 24 seconds. Each trial consisted of 200–250 ms long stimuli (train of 1 ms-long acoustic clicks or electrical/ optical pulses at a rate of 50 Hz in case of predefined oCI/eCI stimuli) repeated at rate of 2 Hz within a response window of 6 seconds. Following perception of a stimulus in the target trial the animal was expected to shuttle (pass over the hurdle) to the opposite side of the ShuttleBox (Figure 3 A). Once the animal shuttled within the response window, the trial was counted as a hit and the stimulus was terminated (Figure 3 A left). Otherwise, an aversive stimulus (mild electrodermal stimulation of the paws via metal bars of the platform, 0.1–1 mA) was executed until the animal shuttled or for maximally 6 seconds and the trial was counted as a miss (Figure 3 A centre). Per session, non-target (no stimulus, no aversive event, Figure 3 A right) and target trials (Figure 3 A left) were randomly intermixed to determine the baseline of the animal activity. The response rate of the animal (Figure 3 D–G) for target/non-target trials is reported as the fraction of target/nontarget trials in which the animal crossed the hurdle divided by the total number of target/non-target trials. In all sessions employing oCI/eCI stimulation each animal was briefly anaesthetised with isoflurane to connect the sound processor and driver circuitry to the implant. Adaptation time was counted once the animal became active again.

After proving that the sound processor can be triggered by acoustic clicks to elicit light stimulation which retrieves pre-trained avoidance behaviour^82^, we moved to the first *in vivo* application of the sound processor using coding strategies. Beforehand we had identified that operating all LEDs at 30 % intensity still was sufficient to trigger behavioural response, pointing towards TL below that value. For MCL estimation we referred to the observation that LEDs operating on 100 % intensity did not cause adverse behaviour in the animals like intensive scratching, which for example had observed for eCI implanted animals. Therefore, for the first implementation of coding strategy we used a TL of −80 dB SPL for 2–8 kHz and −90 dB FS for 8–22 kHz, MCL of −70 dB FS, loudness growth coefficient *L* = 80, and a microphone gain of +12 dB. The processing range of the sound coding strategy was set to the frequency ranges for which aABRs were undetectable, which was 2–20 kHz (Figure 4) and 16–22 kHz (Supplementary Figure 5). Thus, a band width of 18 kHz was coded using 6 LEDs (Figure 4), as those were inside the cochlea, while a band width of 6 kHz was covered by 8 LEDs (Supplementary Figure 5). Within the ShuttleBox sessions either band noise at 80 dB SPL within the predefined frequency ranges (Figure 4) or pure tones at 50 dB SPL (Supplementary Figure 5) within the predefined frequencies were presented as targets. Pure tone of 8 kHz at 80 dB SPL served as control in the rat that still showed residual hearing for this frequency (Supplementary Figure 5 C–E). Plotted response rate to stimuli of a certain band width is the average response rate of the whole session (Figure 4 C and D, 3.4 kHz: 5 frequency bands covering the 2-20 kHz, 4.5 kHz: 4 bands; 6 kHz; 8.5 kHz: 2 bands). To confirm that the observed behaviour triggered by band noise was not cued by residual hearing, we run additional sessions without the oCI system but with the band noise stimuli spanning entire 18 kHz range (2–20 kHz range, Figure 4) or using the identical paradigm as for the session before (pure tones, Supplementary Figure 5 C) in which behaviour was triggered when the oCI system was switched on. Animals did not show the previously learned avoidance behaviour, demonstrating the strict requirement of the oCI system (Figure 4 C and D, Supplementary Figure 5 C).

## Acknowledgements

The authors thank Daniel Weihmüller for PCB layouting, assembly, testing, and help/support in hardware development. We thank Dr. Vladan Rankovic for producing and injecting AAVs. Dr. Daniel Keppeler and Dr. Marcus Jeschke designed the enclosure, which was then produced by Rainer Schürkötter and colleagues (MPI for Biophysical Chemistry, Göttingen). The work was funded by the European Research Council (ERC) under the European Union’s Horizon 2020 research and innovation program (grant agreement No 670759 – advanced grant “OptoHear” to T.M.) and by the Deutsche Forschungsgemeinschaft (DFG, German Research Foundation) via the Leibniz Program (MO896/5 to T.M.) and under Germany’s Excellence Strategy (EXC 2067/1-390729940 to T.M. and EXC 1086 to P.R.). This work was also supported by the Ernst Jung Prize for Medicine (to T.M.).

## Author contributions

T.H., L.J., G.H., and T.M. designed the study. T.H. and L.J. developed software (firmware for sound processor and control software for computer). T.H., G.H., L.J., and T.M. planned the hardware development, which was implemented by G.H. and L.J. supported by Daniel Weihmüller. A.D. and B.W. performed surgeries/implantations of eCIs/oCIs. B.W. deafened the animals. A.D., B.W., and L.J. performed ShuttleBox trainings/experiments involving rats and analysed the data. R.H. designed and generated eCIs. S.A. and P.R. developed and generated oCIs. L.J., T.H., B.W., and T.M. designed the figures. L.J., B.W., and T.H. prepared the figures. A.D. contributed to preparation of the figures. L.J., T.H., B.W., and T.M. prepared the manuscript. All authors discussed the results and commented on the manuscript.

## Supplementary Information

### Choice of the digital signal controller

Prior to starting development of the sound processor, we critically reviewed state-of-the-art DSCs to see which would qualify for the envisioned tasks. The initial benchmarking of DSCs was performed in 2016 and the shortlist included PIC32MZ2048EFH and PIC32MX795F512L (Microchip), KW40Z (Freescale) based on ARM Cortex M0, SAM4L (Atmel) based on ARM Cortex M4, and nRF52832 (Nordic Semiconductor) based on the ARM Cortex M4. A comparison of their key properties is provided in Supplementary Table 1.

We compared and benchmarked the shortlisted systems in various ways. Since the most demanding of the envisioned tasks was the real-time audio processing, we started by bench-marking spectral decomposition methods on all platforms. For that, we implemented spectral decomposition via fast Fourier transform (FFT) and infinite impulse response (IIR) based filter banks in several ways including those optimized for the specific microarchitecture (like the CMSIS DSP library for the ARM-based systems^83^). First, for all filter bank variants we determined the filtering duration of 1 second of audio sampled at 48 kilo-samples per second into 10, 20, and 32 quasi-logarithmically spaced bands followed by an envelope extraction and quantisation in time. This calculation yields a matrix of spectrotemporal data, which can serve as a basis for an *n*-of-*m*-like coding strategy (see Sound coding strategy).

After choosing the ideal implementation for each individual system—as judged by the shortest calculation times for FFT and IIR—we determined the required clock speeds just for real-time spectral decomposition as a function of the number of frequency channels. Finally, based on measurements of electrical current and information from the specifications (combined with data extrapolation where the clock speed was not adjustable as required), we estimated the required power as shown in Supplementary Figure 1. Since CI systems require real-time audio processing for a long period of time, we deem the power efficiency of spectral decomposition to be very important for which nRF52832 (Nordic Semiconductor) outperformed the other processors for more than 20 frequency channels (Supplementary Figure 1).

For the DSCs without built-in radio transceiver we considered various small commercial off-the-shelf Bluetooth Low Energy (BLE) modules to be connected via the Serial Peripheral Interface (SPI) and a few general purpose I/O (GPIO) pins for interrupt signalling. The extra space required by those modules belongs to the downsides of the radioless systems on a chip. First tests with the BLE protocol stack revealed that a sensible amount of working memory must be dedicated to the wireless functionality. Moreover, latencies can become relatively long when operating via BLE, which must be considered in the design of wireless control (for details see Wireless control and communication protocol).

**Supplementary Figure 1.**
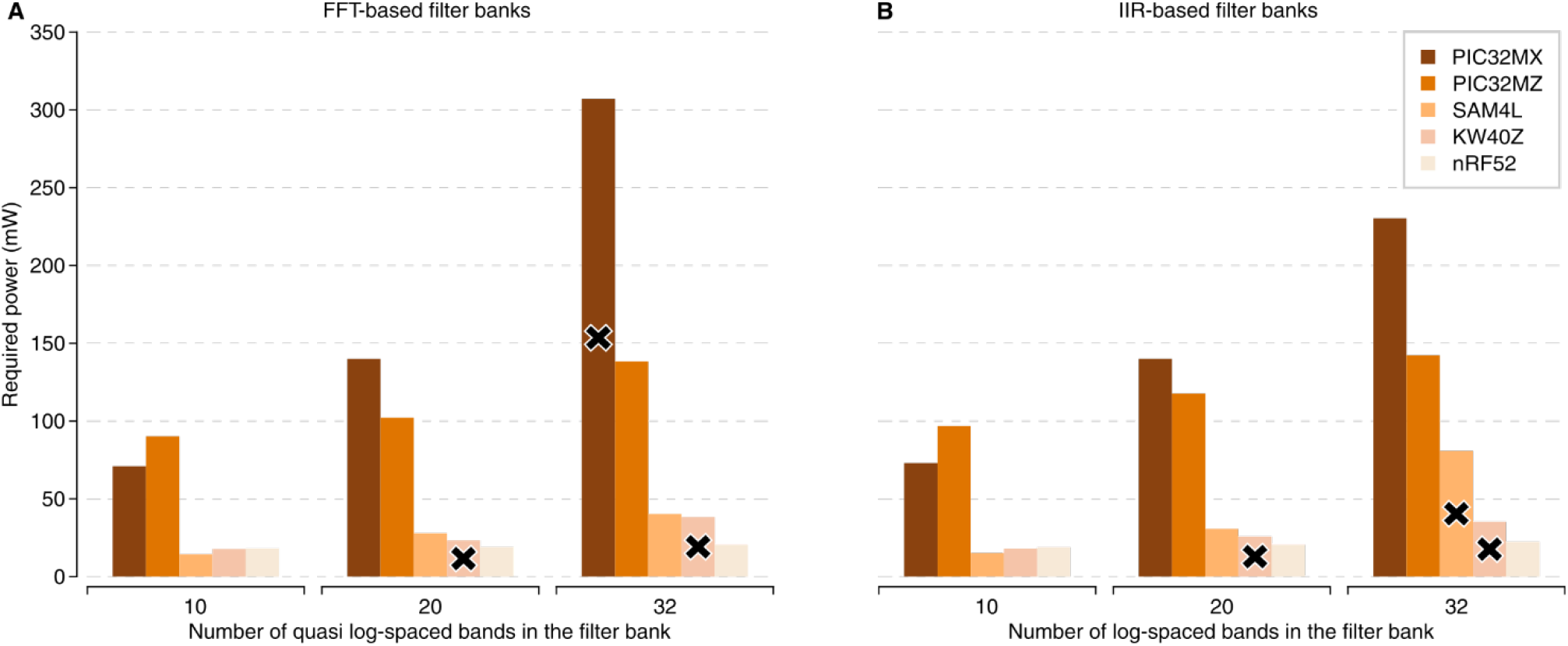
Comparison of power required just for real-time spectral decomposition by digital signal controllers (DSCs). (A) Required power for spectral decomposition via FFT-based filter bank operations. (B) Required power for spectral decomposition via IIR-based filter banks operations. Cross marks denote cases, where the required clock speed for real-time operation was higher than the specified maximum clock speed of the given system.

In a final step, we tested the response speed and quality of the product support, reviewed the clarity and comprehensibility of documentation, as well as the availability of examples and public forums. Eventually, we selected the nRF52832 (Nordic Semiconductor) as implemented in the BL652 (Laird Connectivity) module, which back then, in 2016, was the smallest form factor available on the market that implemented the full reference circuitry as specified by Nordic Semiconductor. While this system on a chip lacks a digital-to-analogue converter (DAC) and the possibility to adjust clock speed, it provides a range of useful interfaces, an on-chip 2.4 GHz radio transceiver, and supports high computing power and efficiency. Last but not least, we found the documentation and support favourable.

### Optical stimulation circuitry

The oCI driver circuity employs a 16-channel digitally-adjustable current sink (TLC5923, Texas Instruments Inc.) and an 8-channel analogue switch (ADG1414, Analog Devices) each controlled by the DSC via dedicated SPIs (SPI0 and SPI1, respectively, see Figure 2). This design enables matrix addressing of the oCI arrays with up to 128 channels: the common anode of the respective LED block (up to maximum 8 blocks) is selectively activated by the analogue switch. The individual LED within a block is selected and its light intensity is adjusted by the current sink, setting of the current level of each LED within a block (up to maximum 16 LEDs per block). Independent of the topology of the LED array, the DSC needs ~15 μs for internal commands to set up the LED array with the requested light intensities. The output of the oCI driver is accessible on the female pin header (CLE-110-01-G-DV, Samtec) for interfacing oCI probes (see Methods). For the future probes employing matrix addressing^32^ pin header with more contacts can be soldered (CLE-114-01-GDV, Samtec).

### Electrical stimulation circuitry

The eCI driver circuity employs a dual 16-channel multiplexer (MAX14661, Maxim Integrated) and a 12-bit DAC (AD5683, Analog Devices) driving a current source (AD8643, Analog Devices) to supply the charge balanced biphasic stimuli via a channel-dedicated capacitor to the selected electrode. The multiplexer and DAC are controlled by the DSC via dedicated SPIs (SPI0 and SPI1, respectively, see Figure 2). By the design of the eCI, the output of the eCI driver is limited to 10 channels accessible on the female pin header (CLE-107-01-G-DV, Samtec) for interfacing eCI probes (see Methods).

### Size, weight, and power consumption

The complete oCI/eCI sound processor and driver circuitry is built in a form of a cylinder accommodated in a low-weight (~6.5 g) and robust plastic enclosure made of Polyether ether ketone (PEEK). The inside diameter of the enclosure is 25 mm and its height 30 mm. The enclosure consists of a threaded head-mounted base (~1.5 g, Figure 1 C and D) fixed by dental acrylic and metal anchors to the skull of the animal, as well as a screwable cap (~5 g, Figure 1 B). This design allows for convenient installation of the device on the animal’s head and battery replacement. The oCI/eCI sound processor and driver circuitry consists of two round multilayer rigid PCBs (Figure 1 B, E, and G), each 20 mm in diameter, carrying commercial off-the-shelf electronic components on both sides, which were interconnected via PCB surface-mounted pin headers (top PCB: FTE-106-03-G-DV, Samtec; bottom PCB: CLE-106-01-G-DV, Samtec). The bottom PCB (closer to the animal’s head) consists of the sound processor and oCI/eCI driver circuitry (Figure 2). It is connected to the head-mounted pin header to interface with the eCI or oCI. The top PCB consists of a battery bracket holding a lithium-ion battery (CP1654A3, VARTA Microbattery GmbH; for details see Supplementary Information), a step-down converter (TPS62240, Texas Instruments Inc.) for powering of the DSC, booster (MCP16251, Microchip Technology Inc.) for powering current source (in case of the eCI system) or LED driver (in case of the oCI system), and a circuit based on a nanopower comparator (MAX9117, Maxim Integrated) that shuts off powering when the battery voltage is too low to allow a reliable operation of the system. A flexible antenna for 2.4 GHz radio (FXP73, Taoglas) is wrapped around the PCBs assembly. The weight of both PCBs including batteries and antenna is ~8 g and complete oCI/eCI system is ~15 g. The performance of the oCI/eCI system operating under battery powering is shown in Supplementary Figure 2.

**Supplementary Table 1.**
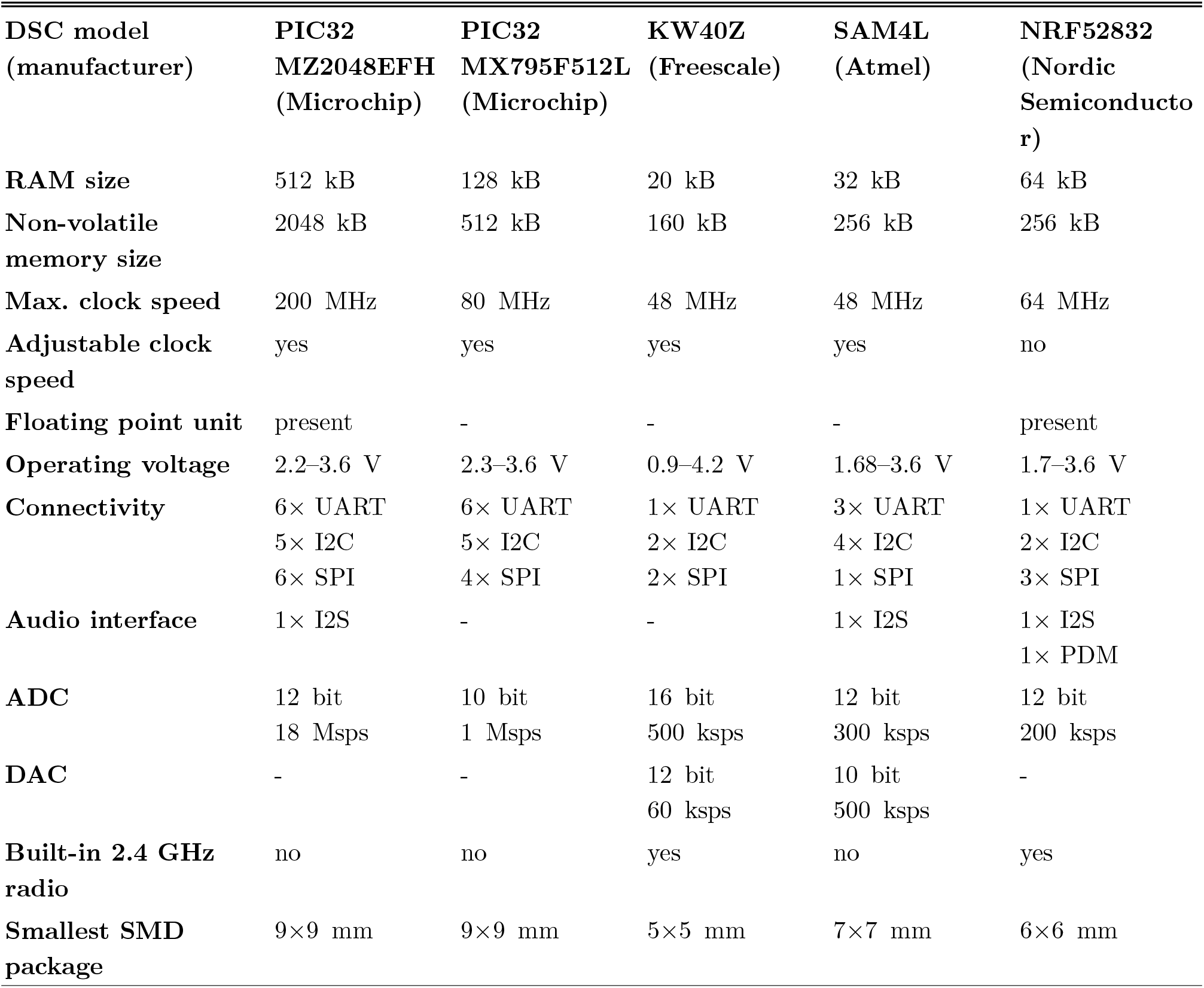
Comparison of the digital signal controller (DSC) specifications. Parameters most instrumental for the DSCs to be used as sound processor and driver for oCI/eCI are presented. There are more than 25-fold differences in random access memory (RAM) and about 12-fold differences in non-volatile memory (NVM) sizes, with more memory meaning more flexibility. The difference in maximum clock speed is less pronounced, and the nR52832 is the only DSC lacking the option to adjust clock speed at all. The range of operating voltage is widest for the KW40Z, while the PIC32 DSCs score with the richest set of interfaces like universal asynchronous receiver-transmitter (UART), inter-integrated circuit (I^2^C) bus, and serial peripheral interface (SPI). Three DSCs feature inter-IC sound (I^2^S) and/or pulse-density modulation (PDM) audio-specific interfaces aiding their use in the sound processing. All DSCs feature a built-in analogue-to-digital converter (ADC; Msps: megasamples per second, ksps: kilosamples per second) required for electrophysiological recordings and battery level monitoring. Three of them lack a digital-to-analogue converter (DAC) needed to precisely adjust the amplitude of electrical stimulation then requiring an external DAC. The two of the five shortlisted DSCs including a radio transceiver happen to have the smallest surface-mounted device (SMD) footprint, which was another key factor for the design. Data obtained from the data sheets provided by the manufacturers.

### Selection of the battery

Seeking the low weight battery enabling reliable operation of the oCI/eCI systems, we tried various batteries starting with lithium coin cell CR1631, including hearing aid dedicated Zinc Air MERCURY-FREE (size p13 and p675, power one VARTA Microbattery GmbH), and cochlear implant dedicated series IMPLANT plus (size p675, power one VARTA Microbattery GmbH). Most of them failed for our purpose not being able to provide enough current for long term wireless operation in permanent receiving or transmitting mode where radio transceiver is always turned on (ShuttleBox paradigm with triggered predefined stimulation, for details see Results and Methods). Finally, we successfully employed a lithium-ion battery (CP1654A3, VARTA Microbattery GmbH). Battery testing running the ShuttleBox-like paradigm (see Behavioural experiments paradigm) revealed 7–8 hours of the device operation. The discharge curve of the battery is shown in Supplementary Figure 2 B.

**Supplementary Figure 2.**
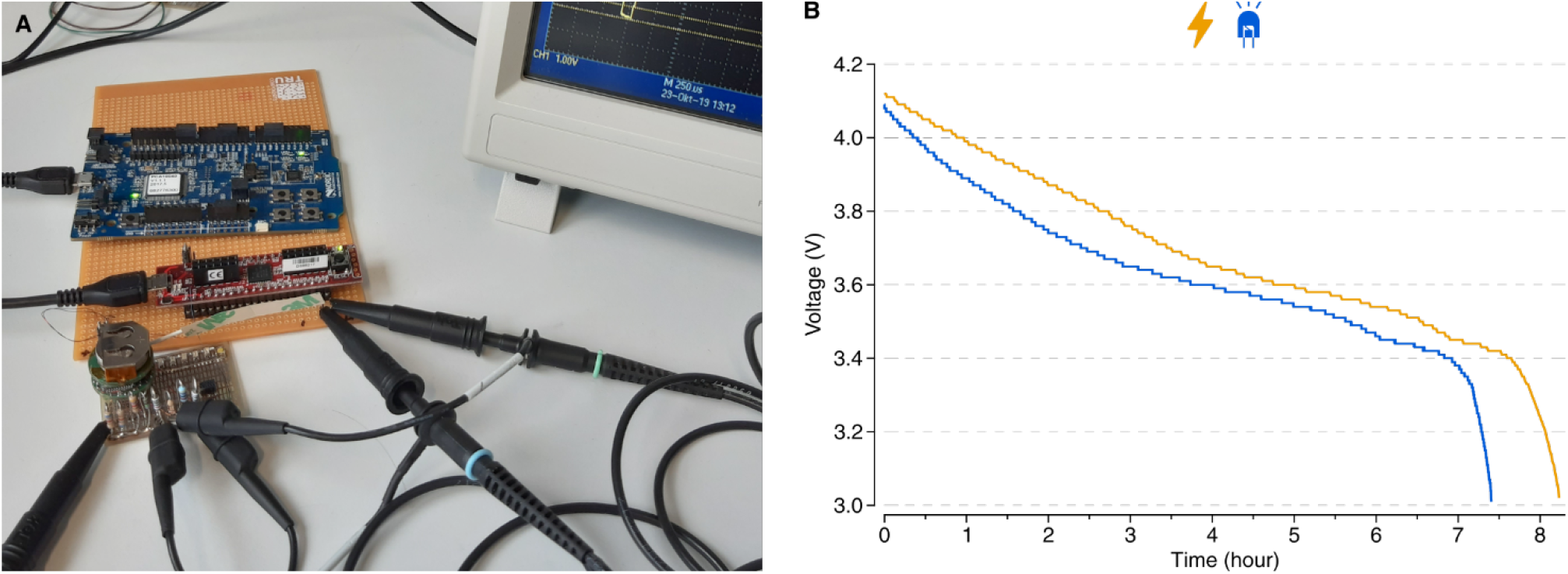
Test of the lithium-ion battery usage under operation in the eCI and oCI systems. (A) Photograph of the custom-build setup employing the most demanding paradigm of the ShuttleBox protocol and a 6.8 kΩ resistor in place of a single eCI channel simulating current injection into the cochlea or single LED of the oCI implant. (B) Corresponding discharge curves of the batteries of eCI (orange) and oCI (blue) systems showing continuous operation for 7–8 hours.

Testing was performed using MATLAB (The MathWorks, Inc) script simulating ShuttleBox experiment in which stimulation is triggered via USB-to-ESB bridge for the maximum length of the response window and minimum inter-trial-interval of 10 s (compare with the ShuttleBox experiment paradigm described in Methods). This simulation is considered the most demanding scenario from the point of the view of power consumption in such ShuttleBox experiments. For eCI system, instead of implant and the animal, the 6.8 kΩ resistor, matching the average impedance of the electrode implanted into the animal, was used. For oCI system, oCI implant was used (see Methods). Stimulus parameters were the same as in the experiments with animals for both systems except of number of channels stimulated simultaneously, here single channel for either eCI or oCI, as well as for eCI current intensity of 300 μA (~20 % of a maximum current possible for an eCI driver) and for oCI light intensity of 20 % of a maximum current possible for an oCI driver (~620 μA per LED). Current consumption of the device and voltage level of the battery were measured with custom-build setup consisting of Cmod CK1 (Digilent Inc.) and a current-shunt monitor INA186 (Texas Instruments Inc.) running custom-coded firmware sending data via UART interface to the PC (Supplementary Figure 2 A).

### Transfer characteristic of the sound processor’s audio system

The audio quality of the sound processor is determined by the acoustic setup (e.g. positioning of the microphone, diameter of the acoustic vent in the PCB, acoustic shadow effects of the processor enclosure), the PDM microphone, and the processor’s PDM interface.

To characterise the overall audio quality of the processor, we developed a program (stored as “Audio calibration mode” in the sound processor, see Figure 2) that analyses signals coming from the PDM microphone and forwards analysis results to a PC. For the analysis it uses a sampling rate of 50 ksps, a 512-point real-valued FFT with floating point precision based on the CMSIS DSP library^80^, and Blackman-Harris window function.

In an acoustically shielded chamber (Industrial Acoustics) we used a single Tannoy Reveal 402 loudspeaker (±3 dB linearity between 56 Hz and 48 kHz) driven by a Zoom UAC-8 audio interface (±1 dB linearity between 20 Hz and 40 kHz at 96 ksps) to play back sound generated by a custom-coded MATLAB (The MathWorks, Inc) script. The distance between the centre of the loudspeaker and the sound processor was exactly 1 m. The sound pressure level was controlled for at the sound processor’s position with a calibrated Phonic PAA3 audio analyser.

Using script, we generated a sine sweep (chirp signal sweeping from 100 Hz to 30 kHz at 96 ksps sampling rate), which we played back at specific sound pressure levels (SPL) while the analysis program was running on the sound processor. The results for 95 dB SPL, 65 dB SPL, and silence (noise floor measurement) for frequencies between 200 Hz and 25 kHz are shown in Supplementary Figure 3.

**Supplementary Figure 3.**
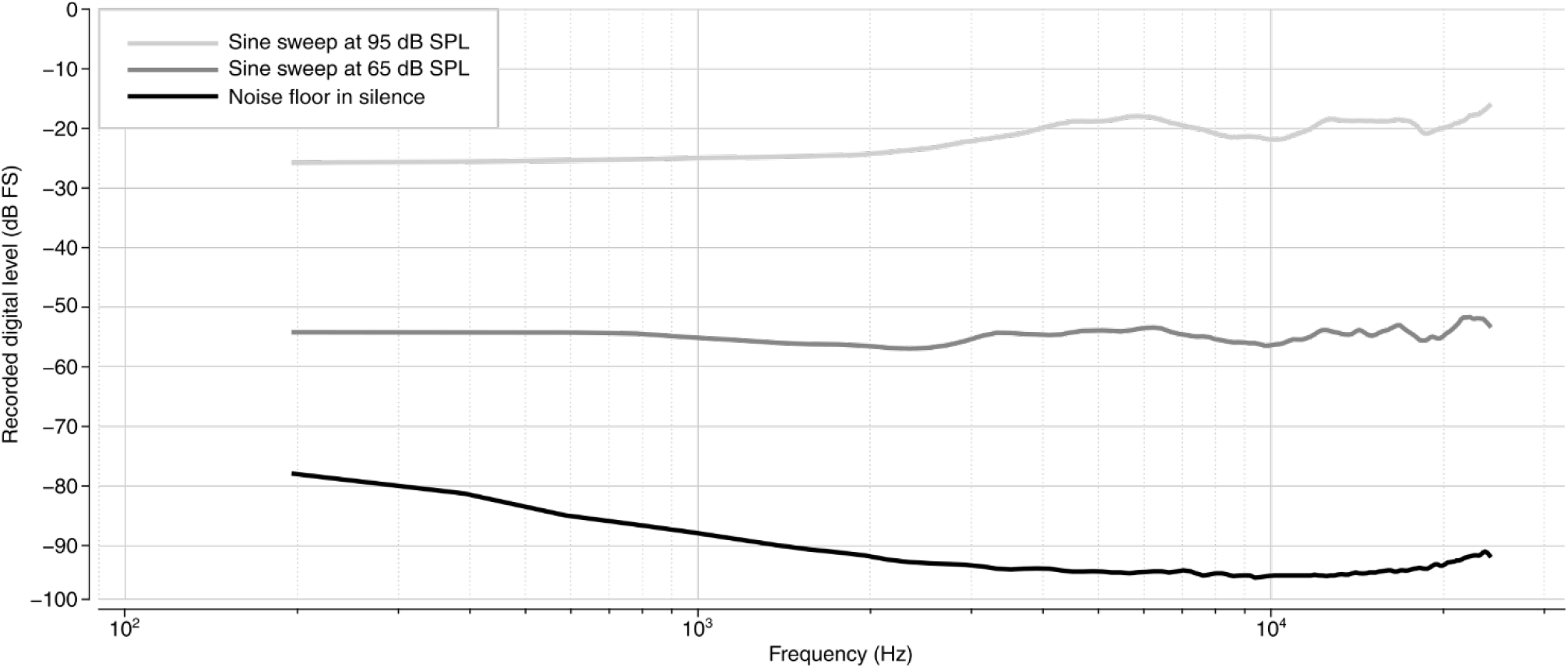
Transfer characteristics of the sound processor’s audio system at different sound pressure levels. Measurements of a noise floor in silence (black) and sine frequency sweep at: 65 dB SPL (light grey) and 95 dB SPL (dark grey).

**Supplementary Figure 4.**
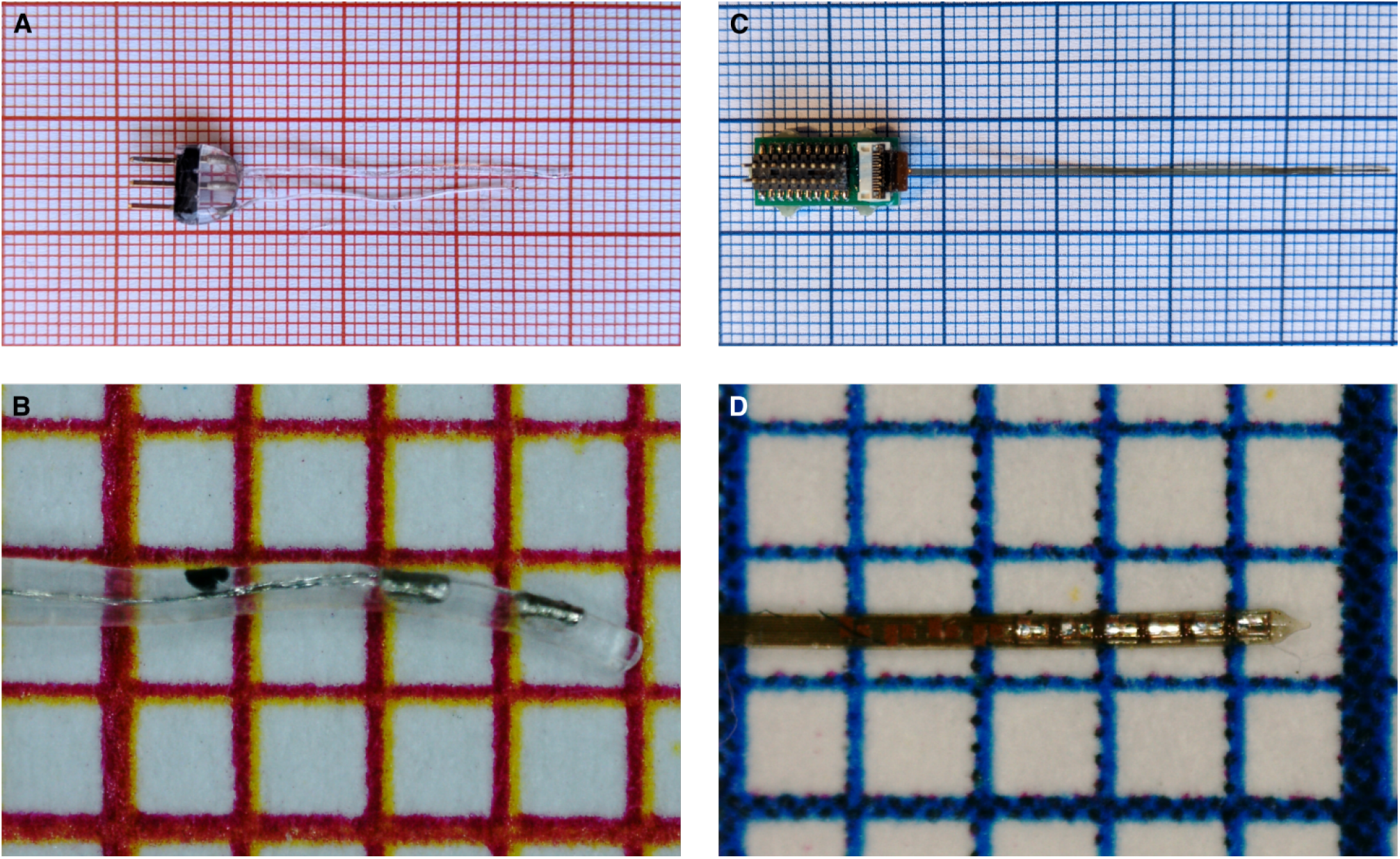
Design of the chronic eCI and oCI probes used in behavioural experiments. (A and B) Photographs of MED-EL’s preclinical two-channel eCI. (A) Two-channel eCI probe showing both intracochlear and return electrodes. (B) Close-up photograph of the intracochlear part with two electrode contacts and black insertion mark of the eCI. (C and D) Photographs of preclinical six-channel oCI. (C) Six-channel LED oCI connected to the adaptor board interfacing ZIF contact pads with the sound processor (left part). The design is identical to the ten-channel probes used in experiment except of lack of four basal LEDs. (D) Close-up photograph of the intracochlear part with six LEDs and contact pads for additional four LEDs. Millimetre paper in the background.

**Supplementary Figure 5.**
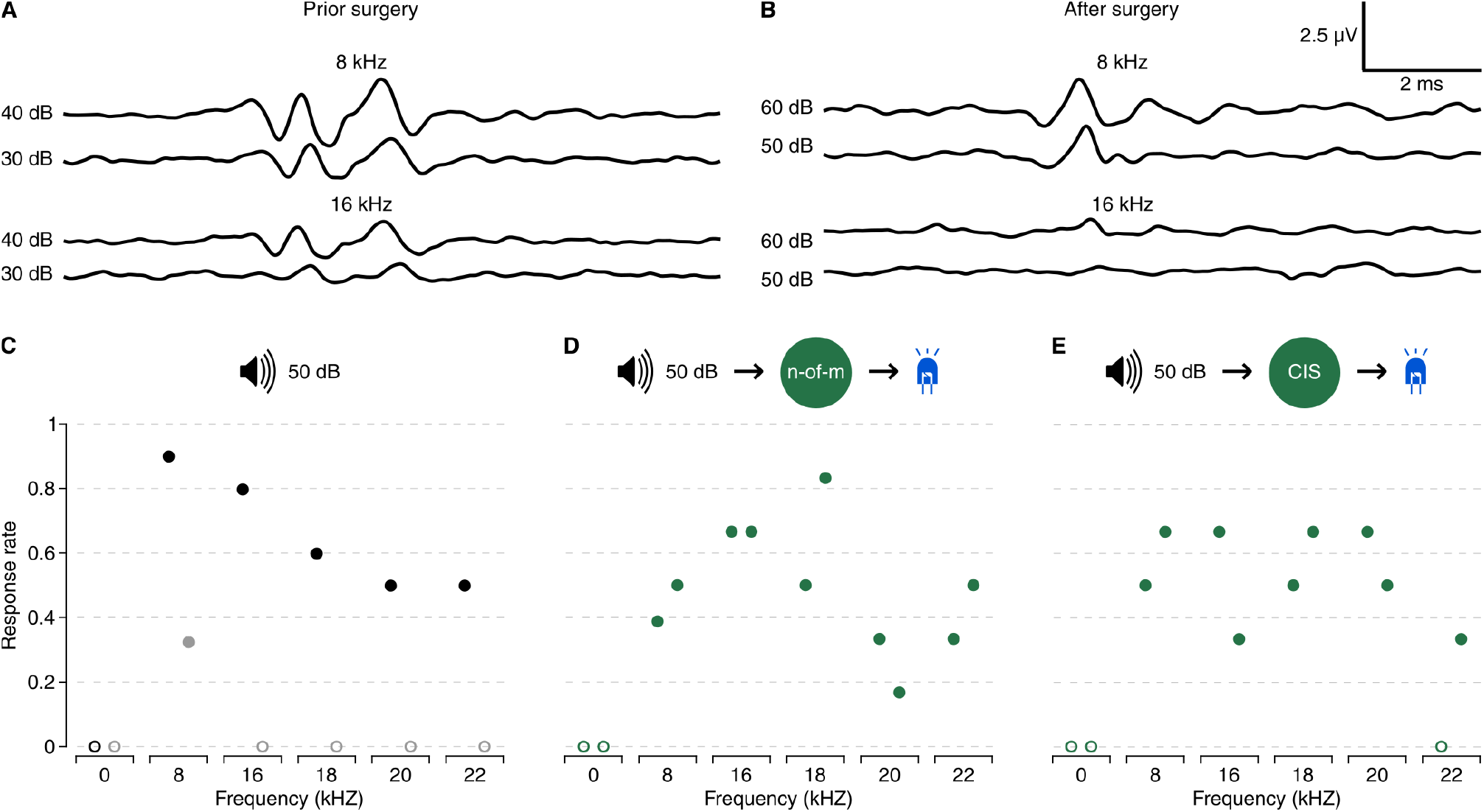
Auditory cued behaviour in a partially deafened rat by an oCI system using different coding strategies of the sound processor. (A and B) Representative aABR, in response to 8 and 16 kHz pulse prior (A) and after (B) the implant surgery and deafening. All traces were recorded from the same animal. (C) Response rate to the acoustic stimulus in the ShuttleBox session of a trained hearing rat (black) and a rat after implant surgery and deafening without the sound processor (grey). (D and E) Results of the ShuttleBox sessions with activated sound processor encoding acoustic stimulus into light stimulus, either using *n*-of-*m* (D) or CIS (E) coding strategy in animals after oCI surgery and deafening (for frequencies ≥ 16 kHz, some hearing was maintained for lower frequencies). The processing range of the sound coding strategy was adapted to the non-audible frequency range of the implanted rat (≥ 16 kHz) and the processor encoded this using 8 LEDs inside the cochlea of the AAV-transduced rat. Filled circles correspond to response rate of one complete ShuttleBox session whereas empty circles emphasize the lack of any response on the behavioural level. While for implanted rat behavioural response was always observed for 8 kHz, a frequency also showing a response in the aABR after surgery (B), behaviour in response to stimulation at 16, 18, 20, and 22 kHz was only observed when the processor was turned on. All acoustic stimuli were presented at 50 dB.

